# Modern human origins: multiregional evolution of autosomes and East Asia origin of Y and mtDNA

**DOI:** 10.1101/101410

**Authors:** Dejian Yuan, Xiaoyun Lei, Yuanyuan Gui, Mingrui Wang, Ye Zhang, Zuobin Zhu, Dapeng Wang, Jun Yu, Shi Huang

## Abstract

The neutral theory has been used as a null model for interpreting nature and produced the Recent Out of Africa model of anatomically modern humans. Recent studies, however, have established that genetic diversities are mostly at maximum saturation levels maintained by selection, therefore challenging the explanatory power of the neutral theory and rendering the present molecular model of human origins untenable. Using improved methods and public data, we have revisited human evolution and found sharing of genetic variations among racial groups to be largely a result of parallel mutations rather than recent common ancestry and admixture as commonly assumed. We derived an age of 1.86-1.92 million years for the first split in modern human populations based on autosomal diversity data. We found evidence of modern Y and mtDNA originating in East Asia and dispersing via hybridization with archaic humans. Analyses of autosomes, Y and mtDNA all suggest that Denisovan and Neanderthal were archaic Africans with Eurasian admixtures and ancestors of South Asia Negritos and Aboriginal Australians. Verifying our model, we found more ancestry of Southern Chinese from Hunan in Africans relative to other East Asian groups examined. These results suggest multiregional evolution of autosomes and replacements of archaic Y and mtDNA by modern ones originating in East Asia, thereby leading to a coherent account of modern human origins.

## Background

There are two competing models of modern human origins termed “Multiregional” and the recent “Out-of-Africa” hypothesis [1]. In the Multiregional model [2–4], recent human evolution is seen as the product of the early and middle Pleistocene radiation of *Homo erectus* from Africa. Thereafter, local differentiation led to the establishment of regional populations which evolved to produce anatomically modern humans (AMH) in different regions of the world, made of four major differentiated groups (Africans, Europeans, East Asians, and Aboriginal Australians). *Homo* has been a single species since the genus first appeared in the fossil record ∼2.3-2.8 million years (myr) ago [2–4]. Support for this model is based on fossils and Paleolithic cultural remains but consistent molecular evidence has been lacking. While autosomal data have put a common ancestor of modern humans at ∼1.5 myr ago, it is still far short of 2 myr [5]. In addition to regional continuity, the model further suggests hybridization among different groups [4]. Seeming difficulties here are the clear separation between modern and archaic mtDNAs and Y, the absence of archaic mtDNAs and Y in modern humans [6, 7], and the young age for the modern Y (∼100 ky) and mtDNA (∼200 ky) [8–10].

The single origin Out of Africa model assumes that there was a relatively recent common ancestral population for *Homo sapiens* which already showed most if not all of the anatomical features shared by present day people. This population originated in Africa ∼200 ky ago, followed by an initiation of African regional differentiation, subsequent radiation from Africa, and final establishment of modern regional characteristics outside Africa [1, 10]. These modern Africans replaced the archaic *Homo* in Eurasia with limited genetic mixing [11–15]. Support for this model comes from the African location of the earliest fossils (∼315 ky ago in Jebel Irhoud, Morocco) of mostly but not all AMH features [16, 17] and the molecular clock or neutral theory interpretation of the greater genetic diversity in Africans [10]. Difficulties with this model include the discrepancy between autosomal and Y/mtDNA age, the Y haplotype A00 with age >300 ky [18], no evidence of first appearance of fully modern humans in Africa, seeming data for multiregionalism within Africa [19] and for multiple dispersals into Asia [20], fossils with AMH features of greater than 85 ky old (upto ∼260 ky) in multiple Eurasia locations (the fully modern teeth in Daoxian Hunan, Longtanshan cave 1 Yunnan, Luna Cave Guangxi, Xuchang Henan, Bijie Guizhou, and Dali Shaanxi in China, Misliya and Shkul/Qafzeh in Israel, and Al Wusta-1 in Arabia) [21–28], and the generally weaker support from fossils and stone tools relative to the multiregional model. While the AMH fossils found outside Africa have been assumed to originate in Africa, an origin in Asia has not been excluded. In fact, in 1983, researchers have derived an mtDNA tree rooted in Asia [29]. Unfortunately, this model was overlooked without anyone ever explaining why the Asia model was less valid than the African Mitochondrial Eve model published 4 years later [10].

Most fatal to the Out of Africa model, however, is that the theoretical foundation for it, the molecular clock and neutral theory, is widely known to be incomplete or has yet to solve the century old riddle of what determines genetic diversity [30]. The neutral theory, while largely sound as a null model and a framework for pre-saturation evolutionary processes, has met with great difficulty as an explanatory framework for most molecular evolutionary phenomena [31–34] and as such should not have been so freely used to account for genetic diversity patterns. Obviously, inferring human origins by using genetic diversity data must wait until one has a complete understanding of what genetic diversity means. The standard for such an understanding should of course be a complete and coherent account of all known puzzles related to genetic diversity.

The unusually admixed features of the Aboriginal Australians have yet to be explained by any model [1]. A list of morphological features aimed at defining modern humans would exclude both modern Aboriginal Australians and Neanderthals, indicating some shared traits between the two [35]. Also unexplained is the origin of Negritos in South Asia. Despite the obvious phenotypic similarities and close Y and mtDNA relationships, no special autosomal relationship has yet been found between Negritos and African pygmies or even among different Negrito groups in South Asia [36].

In recent years, a more complete molecular evolutionary theory, the maximum genetic distance or diversity (MGD) hypothesis, has been making steady progress in solving both evolutionary and contemporary biomedical problems [37–50]. The core concept of the MGD theory, maximum genetic diversity or distance no longer changing with time, is *a priori* expected and supported by numerous facts [37, 51, 52]. In contrast, the neutral theory and its infinite site model fail to take MGD into account and tacitly assume that nearly all observed genetic distances or diversities could still increase with time with no limit defined [53, 54]. The MGD theory has solved the two major puzzles of genetic diversity, the genetic equidistance phenomenon and the much narrower range of genetic diversity relative to the large variation in population size [30, 37]. The primary determinant of genetic diversity (or more precisely MGD) is species physiology [37, 55]. The genetic equidistance result of Margoliash in 1963 is in fact the first and best evidence for MGD rather than linear distance as mis-interpreted by the molecular clock and in turn the neutral theory [37, 39, 44, 45, 56–58]. Two contrasting patterns of the equidistance result have now been recognized, the maximum and the linear [45, 57]. The neutral theory explains only the linear pattern, which however represents only a minority of any genome today. The link between traits/diseases and the amount of SNPs shows an optimum genetic diversity level maintained by selection, thereby providing direct experimental disproof for the neutral assumption for common SNPs [38, 40, 46–48, 50, 59]. More direct functional data invalidating the neutral assumption have also been found [60, 61].

One simple method to determine whether any DNA fragment has reached MGD is to count the number of overlap sites (coincident substitutions) in a sequence alignment of three different species [44]. Such sites represent positions where mutations leading to different residues had occurred independently at the same position in at least two species, which would be a low probability event under the neutral theory or its infinite site assumption but common under the MGD theory [44]. The neutral theory is only valid for slow evolving genes yet to reach MGD, where its infinite site assumption holds and the number of overlap sites follows calculation from probability theory [44]. Unfortunately, however, nearly all existing phylogenetic results are from fast evolving sequences that were *assumed* to follow the infinite site model when they in fact do not as they have now been shown to be enriched with overlap sites [44].

Coincident substitutions at overlap sites do not contribute to genetic distances and make the relationship between distance and time hard if not impossible to model accurately. To overcome this, we have developed the “slow clock” method that only uses slow evolving DNAs with zero or few overlap sites. Contrary to naïve expectations based on the neutral theory, fixed and standing variations in slow evolving proteins are in fact enriched with conservative amino acid substitutions and hence more neutral and suitable for phylogenetic inferences [62]. The method has produced a separation time for the pongids and humans that is remarkably consistent with common sense and the original interpretation of fossil records and drastically different from the result of fast evolving DNAs [45]. Here we used the MGD theory and its related methods to revisit the evolution of modern humans. The unique value of the MGD theory in human origin studies is that it helps select the truly informative sequences that would follow the neutral theory. Once such sequences are selected, the remaining methodologies would be mostly covered by the neutral theory, and we fully grant the neutral theory to be valid for truly neutral sequences still at the linear phase of accumulating variations. We just disagree with its treatments of most genome sequences to be neutral and at the linear or near linear phase of accumulating genetic diversity.

## Methods

### Sequence download

We downloaded ancient and modern human genome sequences using publically available accession numbers. South Asian and Oceanian SNPs data from Pugach et al (2013) were obtained from the authors [63]. The Hunan and Fujian identity information of CHS sample of 1KG were obtained from the Coriell Institute website.

### Selection of SNPs

*Random selection of 255K SNPs as fast evolving SNPs.* We selected 255K SNPs from 1KG data to represent the average variation of the genome (Supplementary Table S1). We first generated a random number for each SNP on a given chromosome followed by sorting the SNPs based on the random numbers, and then selected the top ranked set of SNPs with the number of SNPs in the set proportional to the size of the chromosome. SNPs from the slow set were removed. No consideration for SNP frequency was applied. We used the downloaded VCF files of 1KG to generate the genotyping data.

### Slow evolving SNPs

The identification of slow evolving proteins and their associated SNPs were as previously described [59]. Briefly, to obtain non-synonymous SNPs located in the slowest evolving genes, we collected the whole genome protein data of *Homo sapiens* (version 36.3) and M*acaca mulatta* (version 1) from the NCBI ftp site and then compared the human protein to the monkey protein using local BLASTP program at a cut-off of 1E-10. We only retained one human protein with multiple isoforms and chose the monkey protein with the most significant E-value as the orthologous counterpart of each human protein. The aligned proteins were ranked by percentage identities. Proteins that show the highest identity between human and monkey were considered the slowest evolving (including 423 genes > 304 amino acid in length with 100% identity and 178 genes > 1102 amino acid in length with 99% identity between monkey and human). We downloaded the 1KG phase 3 data and assigned SNP categories using ANNOVAR. We then picked out the nonsyn SNPs located in the slow evolving set of genes from the downloaded VCF files of 1KG (Supplementary Table S2).

### Calling SNPs from genome sequences

We used publically available software SAMTOOLS, GATK, and VCFTOOLS to call SNPs from either downloaded BAM files or BAM files we generated based on downloaded fastq data [64–66].

### Analysis of shared and unique SNPs

Shared and unique SNPs were identified by using downloaded allele frequency information from 1KG. Classification of SNPs into different categories such as intergenic, intron etc were also according to downloaded VCF files of 1KG. For some studies, we also calculated the alternative allele frequency of a SNP in a randomly divided human group by using the PLINK software [67].

### Imputation

Because commonly used SNPs chips for genome wide genotyping have only a fraction of the slow SNPs defined here, we performed imputation to obtain more coverage of the slow SNPs on the South Asian and Oceanian datasets of Pugach et al (2013). We used the SHAPEIT2 software to do phasing for the SNP chip data [68] and the IMPUTE2 software to impute based on 1KG data [69].

### Genetic distance calculation

We used the custom software, dist, to calculate pairwise genetic distance (PGD) or number of SNP mismatches from SNP data [59]. This software is freely available at https://github.com/health1987/dist and has been described in detail in previous publications [41, 70]. We obtained PGD for each of the 25 human groups in the 1KG data and obtained average PGD per group for groups within each of the 5 major continents as represented by the 1KG. We excluded highly admixed groups ASW, ACB, CLM, and PUR in calculating the continental average.

### Principal component analysis (PCA)

PCA is commonly used to discover genetic structures of a population. We utilized GCTA to analyze data in the PLINK binary PED format to generate two files (*.eigenvec and *.eigenva). We drew PCA plot using *.eigenvec file [67, 71]. One sample BEB_HG04131 was found on PC2-PC3 plot to be an outlier and was hence excluded from the PC analysis and most distance calculations presented here.

### Other methods

Other common statistical methods used were Student‟s t test, chi square test, and Fisher‟s exact test, 2 tailed.

## Results

### Contrast between fast and slow evolving DNAs in genetic diversity patterns

Different human groups are well known to share ∼85% of common genetic variations [72]. However, sharing may not necessarily mean genetic exchanges or common ancestry as assumed by the field as saturation or parallel mutations could also explain it. These two explanations could be distinguished by asking whether the fractions of shared SNPs are similarly distributed in the fast versus the slow evolving sequences. Since the majority of human genomes are made of non-coding sequences and hence faster evolving relatively to coding sequences, we randomly selected from the 1000 genomes project phase 3 (1KG) data a set of 255K SNPs to represent the fast evolving SNPs or the average genome wide variation (Supplementary Table S1) [73]. To find the slow evolving SNPs, we first identified the slow evolving proteins by aligning human and Macaca proteomes and then selected only the non-synonymous (nonsyn) SNPs located in these proteins as previously described [59]. Proteins that show the highest identity between human and monkey were considered the slowest evolving, including 423 genes > 304 amino acid in length with 100% identity and 178 genes > 1102 amino acid in length with 99% identity between monkey and human. We downloaded 1KG data and obtained a list of ∼15K nonsyn SNPs located in these slow evolving proteins as our slow set of SNPs (Supplementary Table S2 and S3).

To test the amount of sharing, we examined the SNP frequency files from 1KG. For the three human groups, African (AFR), East Asian (ASN), and European (EUR), we considered a SNP as shared if it has frequency > 0 in more than one group and unique if it is present in only one group. We examined 3 different sets of SNPs, the slow set as defined above, syn SNPs in the slow genes as defined above (Supplementary Table S3), and the random set as defined above. The results showed a clear pattern of more sharing in fast evolving SNPs (Table 1), indicating saturation level of genetic diversity, which further confirmed previous findings of higher genetic diversity in patients of complex diseases relative to normal matched controls [40, 48, 59]. That the observed sharing (24%) in fast SNPs was lower than the 85% for common SNPs was because we did not filter the SNPs by frequency and hence there were many private or low frequency SNPs in our set.

**Table 1.** Sharing of different types of SNPs among three groups. Shared SNPs are present in more than one group and unique SNPs are present in only one group. Shown are fractions of each type of SNPs. SNPs that are not found in any of the three groups (AFR, ASN, and EUR) are grouped as no variations (No var.).

|  | Shared | Unique | No var. | #SNPs |
| --- | --- | --- | --- | --- |
| **Nonsyn slow** | 0.05 | 0.66 | 0.29 | 15422 |
| **Syn slow** | 0.11 | 0.64 | 0.24 | 16591 |
| **Random set** | 0.24 | 0.52 | 0.24 | 254489 |

As fast evolving SNPs are enriched with common SNPs, which may in part explain more sharing as found above, we next studied only rare SNPs with the same allele frequency in a racial group for either fast or slow evolving variants. For each major racial group, we selected a set of SNPs that have the alternative allele appearing only twice in the group (either two heterozygous or one homozygous), as singleton SNPs may be sequencing errors and more common SNPs may be too few for informative studies. We divided the set into several subsets based on evolutionary rates, intergenic, intron, synonymus or missense changes in fast or slow evolving proteins with SNP numbers of a racial group in each subset ranging 281-1376757. Fast evolving proteins here include all those that do not belong to the slow group as defined above. We then examined the fractions of these SNPs that were present in other racial groups. The results showed higher shared fractions for SNPs in fast evolving DNAs except for sharing between AMR (American) and EUR (Fig. 1). As AMR is well known to be highly admixed with Europeans, the observed equal sharing in EUR of SNPs regardless of evolutionary rates in fact made sense and validated our approach. We further validated it by randomly dividing the EAS group into two subgroups EAS1 and EAS2. From the random 255K fast SNPs and the slow SNP set, we selected SNPs that appeared in EAS1 only twice and examined their sharing in EAS2 and other racial groups. The results showed equal sharing of fast and slow evolving SNPs in EAS2 but more sharing of fast evolving SNPs in other racial groups (Fig. 1F). Thus, these results invalidated the present notion of a very recent common ancestor (<100 ky) and large amount of admixture as suggested by the Out of Africa model and support a more distant common ancestor and limited admixture as advocated by the multiregional model.

**Figure 1.**
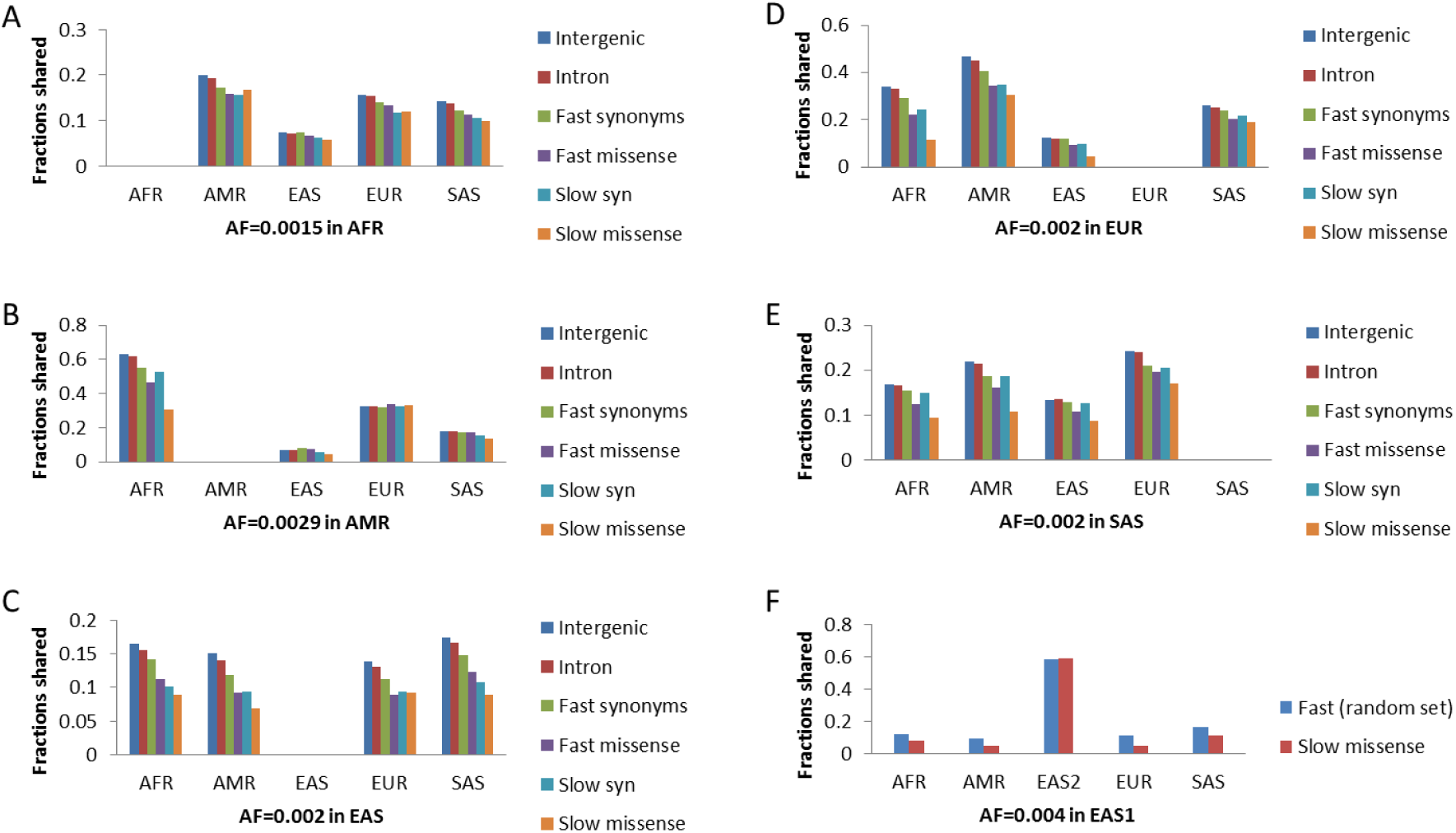
Sharing of SNPs among different racial groups. The fractions of shared SNPs in each racial group are shown for SNPs with alternative allele appearing only twice in AFR (A), AMR (B), EAS (C), EUR (D), SAS (E), or EAS1. EAS1 and EAS2 were two randomly divided subgroups of EAS. These SNPs were classified based on evolutionary rates as indicated.

We next examined the genetic diversity levels within each of the 5 major human groups as sampled by 1KG, AFR, AMR (American), ASN, EUR, and SAS (South Asians), by calculating the average pairwise genetic distance (PGD) per group in different types of SNPs, including the slow set and the random set as defined above, and a stop codon gain/loss set (Fig. 2). In our analysis here, we have excluded 4 highly admixed groups, Americans of African Ancestry in SW USA (ASW), African Caribbeans in Barbados (ACB), Colombians from Medellin Colombia (CLM), and Puerto Ricans from Puerto Rico (PUR). Since certain deleterious SNPs may exist only in heterozygous (het) state rather than homozygous (hom) state, we calculated, in addition to total PGD contributed by both het and hom differences, also the hom PGD resulting from hom mismatches that should better represent neutral diversity. As shown in Fig. 2, hom PGD showed different pattern from total PGD only in the slow SNPs, with the hom PGD level of AFR below the average of five groups while that of AMR being the highest. Remarkably, the stop codon set showed similar pattern as the random set, with AFR having the largest PGD. This indicates functionality rather than neutrality for the random set since stop codon SNPs are definitely functional given its dramatic effect on protein structure [61] and since a neutral set is expected to be very different from the stop codon set. To verify the results of stop codon SNPs, we also found similar PGD pattern in a set of splicing site gain/loss SNPs that are also expected to be functional (Supplementary Information S1 and Fig. S1A-B). Overall, these results showed Europeans with the lowest diversity in stop codon and splicing SNPs and East Asians with the lowest diversity in the random set (P<0.01). Africans have the highest genetic diversity levels in all types of non-neutral SNPs examined as well as the random set (P<0.01), thereby deeming the Out of Africa model untenable as it is based on the now disapproved assumption of neutrality for a random set of genome wide SNPs.

**Figure 2.**
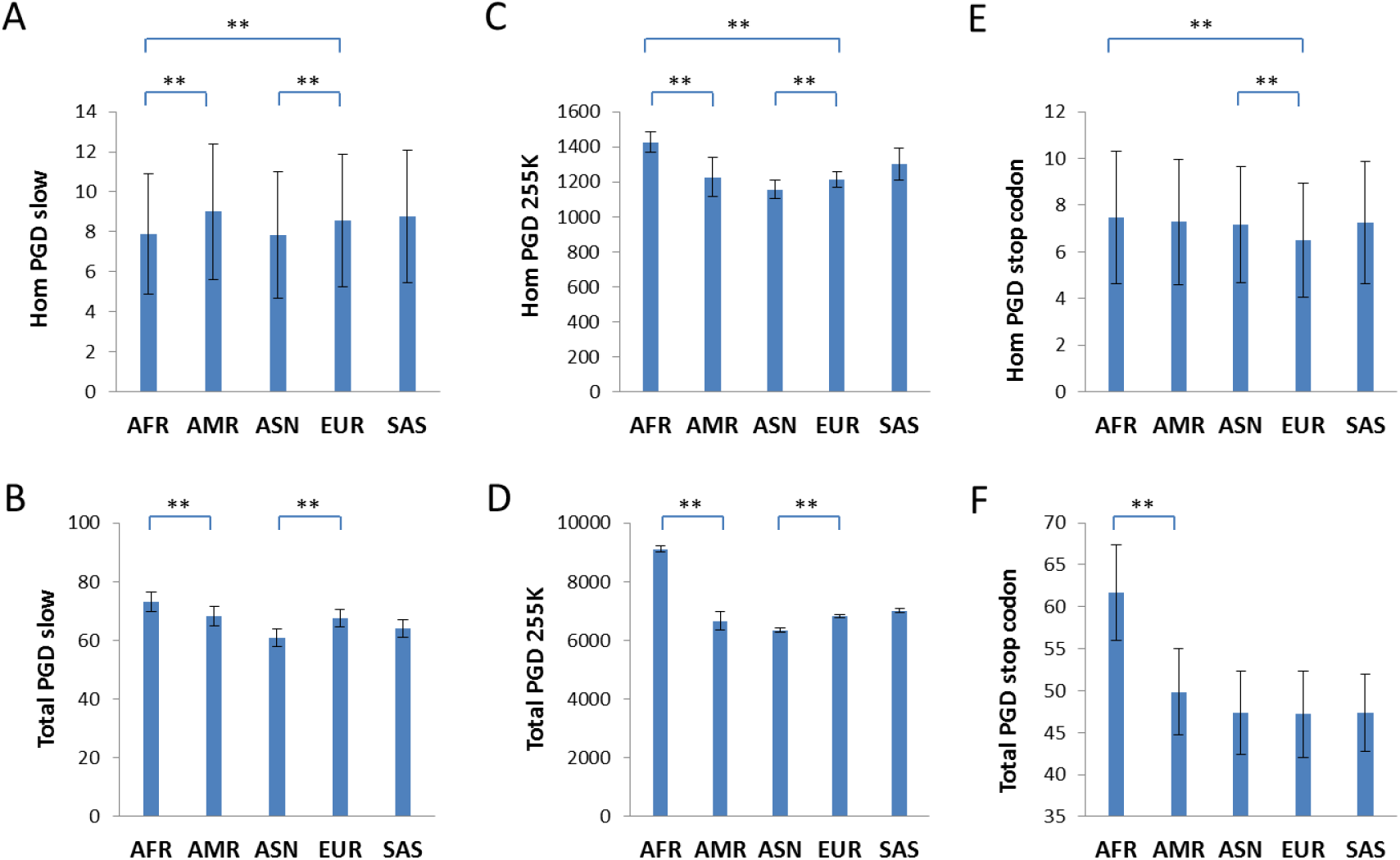
Pairwise genetic distance as measured by different types of SNPs. Pairwise genetic distance (PGD), either by homozygous mismatches (Hom) or by both homozygous and heterozygous mismatches (Total), as measured by three different types of SNPs is shown for each of the 5 major human groups in the 1KG. Known heavily admixed groups such as ASW and ACB in the African group or CLM and PUR in the American group were excluded in the analysis. Data are means with standard deviation. **, P <0.01, t test, 2 tailed.

To confirm if we have made the appropriate cut-off in selecting the slow SNPs as our phylogeny-informative set of neutral SNPs, we verified that the next set of just slightly less conserved nonsyn SNPs (total number ∼13.7K, Supplementary Table S4) within 361 autosomal proteins already behaved like the random set or the stop codon set (800-1102 aa in length with identity between human and monkey >99% but <100%) (Supplementary Information S1, Fig. S1 C-D). Furthermore, syn SNPs within the slow set of proteins as defined above (Supplementary Table S3) gave PGD patterns similar to the stop codon SNPs but unlike the nonsyn SNPs within the same set of proteins (Supplementary Information S1, Fig. S1 E-F). Finally, we confirmed that these slow evolving proteins still have neutral nonsyn variations that are not under natural selection by showing that these proteins have fewer overlap or recurrent mutation sites than relatively faster evolving proteins (Supplementary Information S2 and Table S5), and that known positively selected genes are faster evolving (Supplementary Information S3). Together, these results suggest that only hom distance calculated from the slow nonsyn SNPs, hereafter referred as the slow SNPs, can be informative to phylogenetic inferences. To view slow nonsyn SNPs as neutral seems counter to expectations based on the neutral theory but in fact makes sense under the MGD perspective as fast SNPs are at saturation levels that are maintained by selection and play adaptive roles in response to fast changing environments [62].

Using hom distance measured by slow SNPs, we found, as expected, Africans as the outgroup to the other 4 groups as sampled in 1KG because the non-African groups are closer to each other than to Africans (Supplementary Fig. S2A). Also as expected from *a priori* reasoning but not from the existing model, Africans are closer to each other than to non-Africans. However, for the random set of 255K SNPs, total distance within Africans was similar to that between Africans and non-Africans, which is well known from previous studies and reflects saturation as we now realized from the MGD theory (Supplementary Fig. S2B). This result also established the maximum genetic equidistance phenomenon, previously known only at the inter-species level, at the intra-species level where groups with lower MGD are equidistant to the group with the highest MGD with the distance being equal to the MGD of the highest MGD group. The result independently confirms the difference between slow and fast SNPs and the fact that fast SNPs are at saturation level of genetic diversity.

We also reexamined the claim of an inverse correlation between intra-population genetic diversity and distance to Africa, purporting to support the Out of Africa model [74–76]. Clearly, such correlation was not observed for the slow SNPs where Americans have the highest diversity among all groups (Fig. 2). Neither was it found for the 255K fast SNPs where Americans were more diverse or have higher PGD and higher numbers of het alleles than East Asians; South Europeans (IBS and TSI) were more diverse than North Europeans (CEU and FIN); South Asians were more diverse than Europeans (Supplementary Fig. S3). Different selection pressures on the fast SNPs for different populations may explain these patterns. Ascertainment bias plus overlooking saturation may explain previous conclusions.

### Divergence time between major human groups

To estimate the time of separation between major human groups, we determined the mutation rate of the slow evolving genes. We found 34 informative genes in the 178 slow evolving genes as defined above that showed gap-less alignment in any pair of comparisons among humans, chimpanzees, orangutans, and monkeys (Supplementary Table S6). Assuming gorilla and orangutan contributed similarly to their genetic distance since their split 12 myr as inferred from the fossil records [77], we obtained a gorilla or orangutan mutation rate of 0.000173 aa per myr per aa for the 34 genes (47628 aa). Given a distance of 0.00385 aa per aa between human and orangutan and their separation time of 17.6 myr [45], we used the formula 0.00385 = R_human_ × 17.6 + 0.000173 × 17.6 to obtain the human mutation rate as 4.57E-5 aa per myr per aa, which is 3.88 times slower than orangutan‟s. Given this mutation rate and the distance matrix (total distance including both het and hom distances) as shown in Table 2 (only the largest distance among groups are shown), we estimated the split time between ESN (Esen in Nigeria) and GBR (British in England and Scotland) as 1.92 myr, consistent with the known first migration out of Africa for the Homo species as shown by the fossil records. The split between ESN and CHS (Southern Han Chinese) was similar or slightly shorter at 1.86 myr and not significantly different from that between ESN and GBR. In fact, using hom distance as measured by the slow SNPs which represent neutral distance better, ESN is slightly closer to CHS (14.87) than to GBR (14.93). We only used the largest distance between groups, which was between ESN and GBR, to calculate the time in order to be more precise. Since admixture was common, shorter distances between some pairs of groups may be a result of gene flow and hence not reflect true separation time.

**Table 2.**
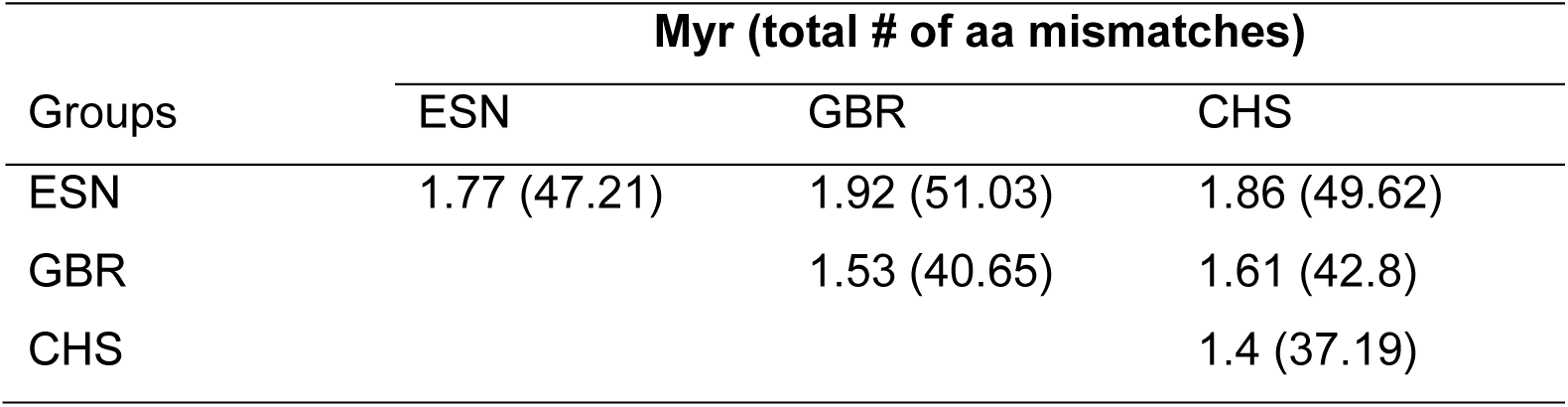
Time of divergence between human populations. The separation time and average pairwise genetic distance (total distance including both het and hom distances) between human populations (ESN, GBR, CHS) in 9578 slow evolving autosome SNPs located in the 178 genes (>99% and <100% identity between human and Macca) with total length 291083 aa. The human mutation rate was estimated as 4.57E-5 aa/myr/aa × 291083 aa = 13.32 aa/myr.

| Groups | Myr (total # of aa mismatches) |  |  |
| --- | --- | --- | --- |
|  | ESN | GBR | CHS |
| ESN | 1.77 (47.21) | 1.92 (51.03) | 1.86 (49.62) |
| GBR |  | 1.53 (40.65) | 1.61 (42.8) |
| CHS |  |  | 1.4 (37.19) |

### Y chromosome phylogeny

The existing Y phylogenetic tree depends on inferring derived alleles and in turn requires the validity of the infinite site assumption, which means no maximum genetic distance and no recurrent mutations. However, this assumption can be proven invalid even just by the existing Y tree itself, since the tree shows numerous recurrent mutations that were simply ignored without valid reasons (Supplementary Table S7), especially for the early branches with some such as KxLT and HIJK contradicted by as many as 50% of all relevant SNPs [78]. That these self-contradictions mostly occurred for the early African branches such as BT and CT but rarely for the terminal Eurasian ones indicates the unrealistic nature of these early branches. Also, while haplotypes with few sequence variations from the ancestor of F, C, D, E, NO, KxLT, or K are routinely found in present day people, none could be found for these early branches. The branching pattern in Africans often involves one branch, such as A00, with few or no sub-branches while the other branch A0-T accounting for all of the remaining haplotypes on Earth, which is odd and against branching patterns known in experimental biology such as the embryonic differentiation into three layers with each layer giving rise to multiple cell types.

Given functionality for genome wide autosomal SNPs as discussed above, it is easily inferred that most SNPs in Y chr are also non-neutral. Evidence for extreme natural selection on Y is also known [79, 80]. We therefore redrew the Y tree based on shared alleles (rather than on derived alleles), which may mean common physiology more than common adaptations if physiology is the chief determinant of MGD. Using previously defined haplotypes for 1KG samples (Supplementary Table S8) and 58251 cleanly called SNPs (no individual with uncalled SNPs, Supplementary Table S9) [81], we found a major megahaplogroup ABCDE (Fig. 3). Megahaplotype F, defined as lacking any mutations that define other haplotypes, is the ancestor. All F-like or F* haplotypes sequenced so far are partial ABCDE carrying 4 (Lahu_HGDP01320), 13 (Malay_SSM072), or 14 (KHV_HG02040) of the 151 mutations that group ABCDE (Fig. 3) [81–83]. The F* haplotype is most common in East Asia, present in 5 of 7 (71.4%) Lahu males in Yunnan of South West China [84], 10-15% of Han and other minority Chinese, and low percentages (<10%) in South Asians and French. Furthermore, the top 4 individuals among 1KG closest to the ∼45 ky old Western Siberian Ust’-Ishim who carried NO haplotype and was expected to be most like the AMH ancestor were all East Asians with Asian haplotypes F and O (F2 in KHV_HG02040, O2 in CHB_NA18534, O3 in CHS_HG00559, O3 in KHV_HG02088), indicating least deviation from the ancestor for Asian haplotypes [14]. Those three O type East Asian individuals also were the closest to the three F* carrying individuals above. These results suggest the origin of F in East Asia with subsequent migration to other regions of the world (Supplementary Fig. S4).

**Figure 3.**
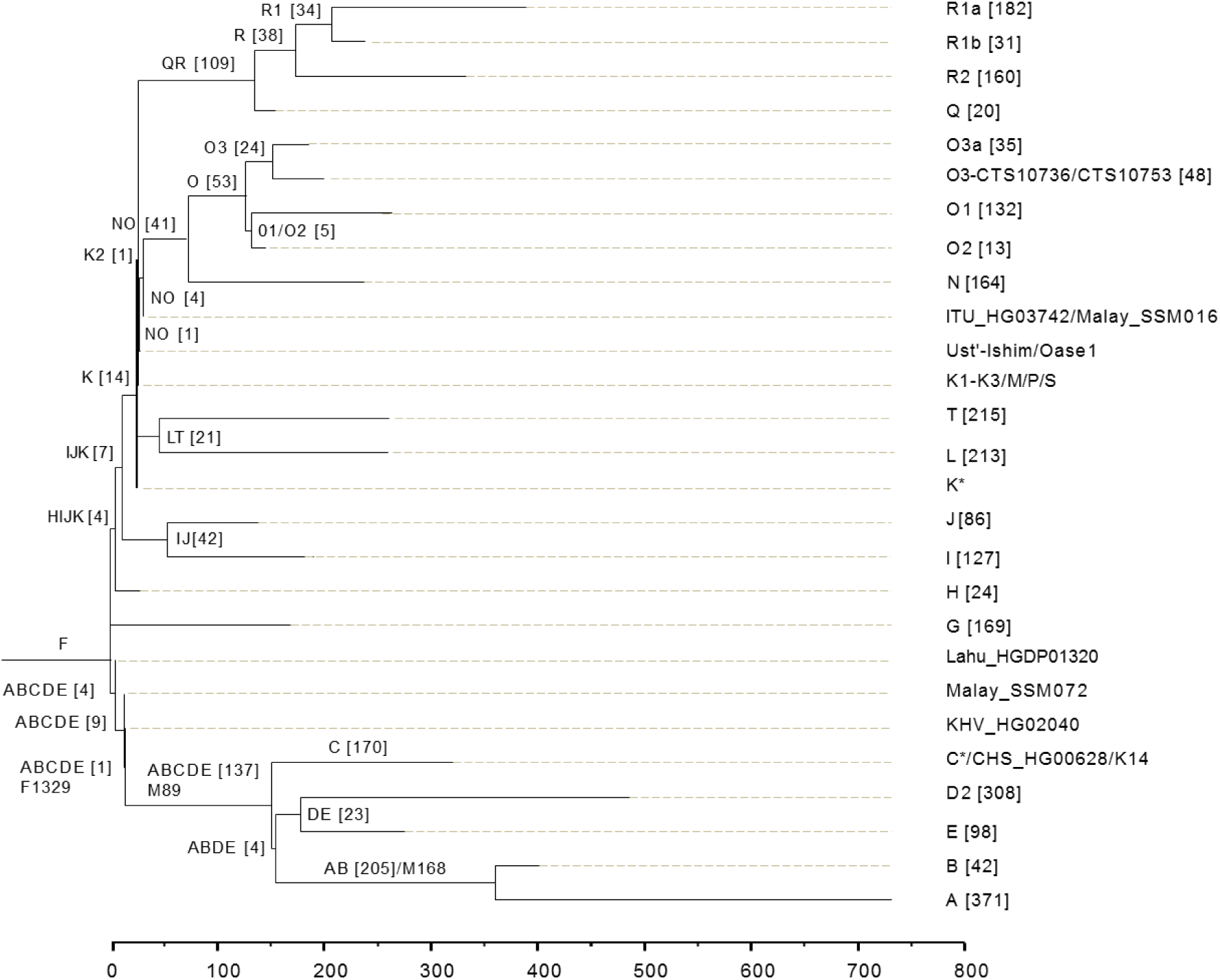
Y chromosome phylogeny. Branch lengths are drawn proportional to the number of SNPs. Only major haplogroups are shown with defining SNPs indicated for some. Numbers in parenthesis indicate the number of SNPs defining a haplogroup among the 58251 cleanly called SNPs in the 1KG. Individuals with few changes from an ancestor haplotype are also listed as shown.

In our tree, alleles previously used to define BT and CT now merely represent alleles associated with the original F ancestor. The AB grouping in our case makes more sense with phenotypes than the BT grouping since it groups African B with African A rather than with the CT group containing mostly Eurasians. The key feature of our tree is that every haplotype besides the original F is associated with haplotype specific SNPs and there are no inconsistent SNPs. Such self-consistency alone would qualify it as more correct than the self-inconsistent tree rooted in Africa.

A real haplotype should exist in a way that has only its own haplotype specific SNPs plus private SNPs. While it is more likely to be the case for terminal haplotypes, it is not impossible for ancestral haplotypes close to the root of the tree, which could be used to distinguish two different competing classifications on an ancestral haplotype. One of the major differences between our Out of Asia tree and the Out of Africa tree is the position of haplotype C, which belongs to ABCDE in the former and CT in the latter. The ABCDE haplotype is closer to the root in the Asia tree (among the first to branch out from the root) than CT is in the Africa tree (2^nd^ to branch out from the root). Hence, relative to ABCDE, CT should have a higher chance to be like a more terminal haplotype. The number of CT defining SNPs is larger than ABCDE (264 vs 151, with additional 50 SNPs contradicting CT), which should also make CT more like a terminal branch. In reality, however, one found the exact opposite. People with only ABCDE specific SNPs plus private SNPs have been found as in present day people Lahu_HGDP01320, Malay_SSM072, and KHV_HG02040 (Fig. 3). That these Y chr had only a portion of ABCDE defining SNPs is consistent with the dynamics of the ancient appearance of ABCDE. Also consistent with the ancestral status of ABCDE, the 38.7 ky old Kostenki14 had 83 among 84 informative SNPs supporting it as ABCDE and 88/92 as C, and the 30.6 ky Vestonice43 had 19/20 as ABCDE and 20/22 as C [85, 86]. In contrast, no one with only CT specific alleles plus private SNPs has been found to exist today or in the past. All ancient Europeans known to be CT and lacking alleles for downstream haplotypes were in fact missing informative sites for at least some haplotypes due to incomplete coverage in sequencing [85, 86]. These findings strongly support grouping C in ABCDE rather than in CT, thereby invaliding the Out of Africa tree.

### mtDNA phylogeny

The existing mtDNA phylogenetic tree has exactly the same problems as the existing Y tree as discussed above. mtDNA has also been found to be under strong selection relative to more slowly evolving nuclear DNAs, consistent with the MGD theory [34, 87, 88]. Based on previously defined mtDNA haplotypes for 1KG (Supplementary Table S8)[81], we redrew the mtDNA tree using slow evolving SNPs, which alter amino acids or RNA sequences (Fig. 4A, Supplementary Information S4, Supplementary Fig. S5, Supplementary Table S10). Fast SNPs are more involved in adaptation to fast changing environments and should not be used whenever possible. Two lines of evidence suggest haplogroup R as the ancestor of all modern haplogroups. First, ancient humans are expected to be closer to the ancestor and the oldest AMH, Ust’-Ishim, carried the R* haplotype [14]. Second, R0 is the least differentiated haplotype and closest to the ancient haplotype in Ust’-Ishim (Fig. 4B). That R0 is most common in Chinese among 1KG samples indicates origin of R in East Asia (Fig. 4B) and subsequent diversification in other regions of the world (Supplementary Fig. S6).

**Figure 4.**
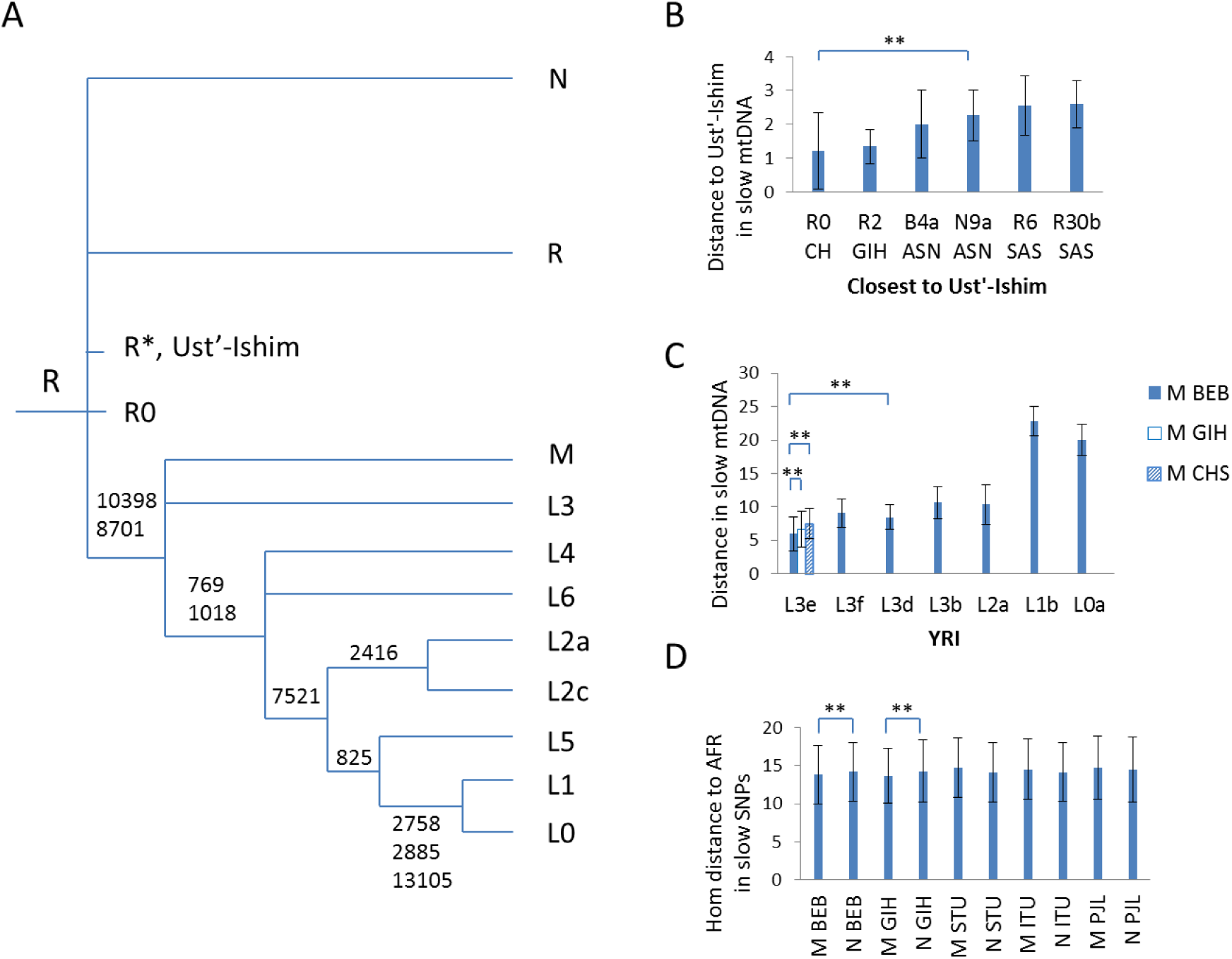
mtDNA phylogeny. (**a**) The mtDNA tree was drawn using slow evolving SNPs as indicated with the common ancestor haplotype defined as being closest to the ∼45,000 year old Ust’-Ishim. Only major branches are shown and no slow SNPs could be found to separate N and R. (**b**) Genetic distance in slow mtDNA SNPs to Ust’-Ishim mtDNA for haplotypes in 1KG. Only the closest few are shown. (**c**) Genetic distance in slow mtDNA SNPs to the M haplotype in BEB, GIH, or CHS for different L haplotypes in the YRI group.(**d**) Genetic distance in slow autosomal SNPs to individuals of South Asian BEB (or GIH, STU, ITU, PJL) carrying either the M or N haplotype. Data are means with standard deviation. **, P<0.01, t test, 2 tailed.

Unlike Y, mtDNA diversification as defined by slow SNPs here is far more star like with multiple parallel haplotypes and few hierarchical structures (Fig. 4A, Supplementary Fig. S5), which is expected from the vast difference in the possible number of offspring between males and females. Many female individuals with R0 might each serve as an ancestor of a specific haplotype within R or N haplogroup, and R is not a sub-branch of N. M also directly derived from R0. L and numerous M subtypes shared a few defining SNPs.

To confirm M giving rise to L, we examined mtDNA distance between African (YRI: Yoruba in Ibadan, Nigeria) L and South Asian (BEB: Bengali from Bangladesh) M and found L3e to be the closest to M (Fig. 4C). Also, M of BEB or GIH is closer to L3e than M of CHS, indicating a more direct role for BEB or GIH in dispersing AMH mtDNA into Africa and a Southern route into Africa. Consistently, in autosome distance, BEB or GIH with M haplotype were closer to Africans than those with N (including R) haplotype (Fig. 4D), despite the fact that people with M had larger autosomal nucleotide diversity than those with N (PGD: M_BEB = 8.59, N_BEB = 7.9, M_GIH = 8.42, N_GIH = 8.36). M may be closer to the common ancestor of M and L since all L types except L3 are at least 2 fixed slow SNPs (769, 1018) away from the common ancestor. There might be a time when there were multiple M types with no L, and then one of the M types with mutations in 769 and 1018 sites became the common ancestor of all L types except L3.

### Neanderthals and Denisovans

If major human groups have separated ∼2 myr ago with region specific features developed not long after separation such as shovel shaped teeth in *H. erectus* from China (Yuanmou man and Peking man), Neanderthals and Denisovans with features more modern than *H. erectus* should be expected to belong to one of the modern groups today. However, previous studies have found Neanderthals to be outgroup to AMH and used D-statistics to show Neanderthal gene flow into non-Africans but oddly not Africans [11, 12]. The assumption of D-statistics is that all modern groups are equidistant to chimpanzees so that presence of derived alleles (different from chimpanzees) was due to gene flow from Neanderthal. If in fact Africans are closer to chimpanzees or carrying more ancestral alleles in general due to perhaps common adaptation to the Africa environment, the conclusion of gene flow into non-Africans would become invalid. We examined this by measuring genetic distance between 1KG and 10 previously sequenced chimpanzee genomes [89]. Using the random 255K SNPs set, we found closer hom distance between Africans and chimpanzees than between non-Africans and chimpanzees (Supplementary Fig. S7). As presence of Neanderthal derived alleles in a non-African are mostly in het state [14], which could be observed to be biased toward non-Africans only if Africans are in hom ancestral state, the fact of more hom ancestral alleles in Africans therefore deems invalid the previous finding of Neanderthal gene flow into non-Africans. Furthermore, as already noted above for Y and mtDNA trees, the finding of saturated level of genetic diversity makes the infinite site assumption invalid, which in turn makes the assignment of ancestral and derived alleles unrealistic. That the D statistics method may not be appropriate to detect Neanderthal introgression has also been independently found by others [90]. Thus, the relationship between Neanderthals/Denisovans and present day populations remains to be determined.

Making use of the published Neanderthal and Denisovan genomes [11, 12, 91], we calculated the genetic distance in slow SNPs between 1KG and Neanderthals (Altai, Vindija 33.16, 33.25, 33.26, and Mezmaiskaya1) or Denisovan3 (Fig. 5A). These ancient genomes showed closer distance to Africans except Vi33.25 to ASN and Vi33.26 to AMR. Denisovan3 was closer to Africans than Neanderthals were (Fig. 5A). In contrast to the Neanderthals and Denisovan3, their near contemporary AMH Ust’-Ishim from Western Siberia was closest to SAS (Fig. 5A) and grouped with SAS in PCA plots (4B-G). We also studied the more recently reported Neanderthal genomes of Vi33.19, Vi87, Les Z4, GoyetQ56, and Mezmaiskaya2 [6, 92] and found them to be also closest to AFR except Vi87 who was equally related to AFR and SAS (Supplementary Fig. S8). Using the slow SNPs but not the non-informative SNPs, we also found that two Neanderthals from the same location, Mezmaiskaya 1 and 2, but separated for ∼20 ky were in fact closest to each other than either was to any other Neanderthals or ancient DNAs of modern humans found anywhere in the world (Supplementary Fig. S9), thus confirming regional continuity and invalidating the previous conclusion of Neanderthal population turnovers [6, 92]. These results suggest that Neanderthals and Denisovans were Africans who migrated into Eurasia and admixed with local non-Africans. The observations of an East Asian like Neanderthal (Vi33.25) in Europe at >45,000 years ago and of a South Asian like Western Siberian (Ust’-Ishim) from ∼45,000 years ago indicates migration of Asians into Europe around the time of AMH origin in South East Asia.

**Figure 5.**
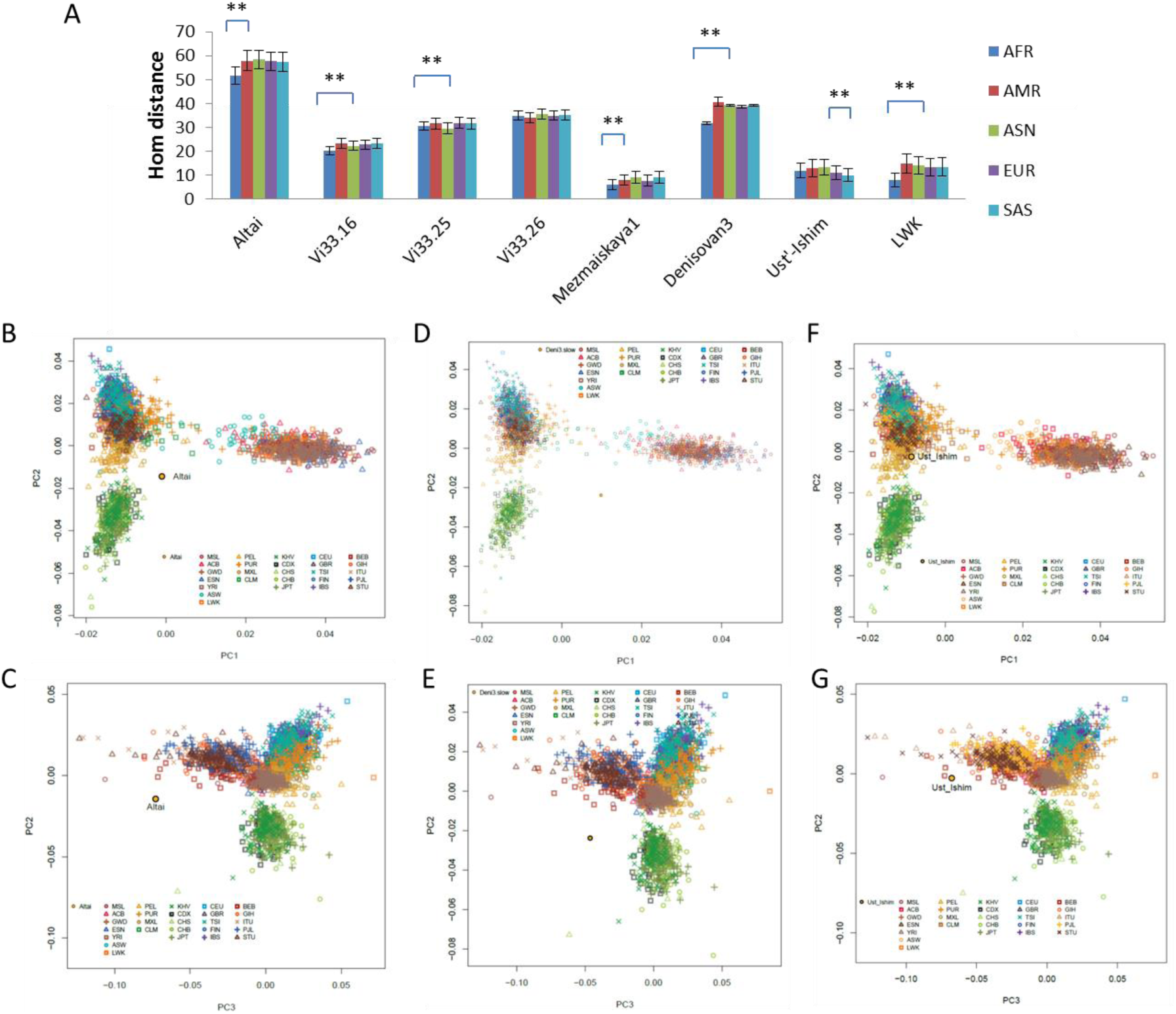
Autosomal relationship between archaic and modern humans. (**a**) Shown are the genetic distances between the 5 groups of 1KG and Neanderthals, Denisovan3, Ust’-Ishim, or the modern African group LWK. Data are means with standard deviation. (**b-g**) PCA plot analyses (b, d, f for PC1-PC2; c, e, g for PC3-PC2) for Altai, Denisovan3, or Ust’-Ishim merged with 1KG. **, P<0.01, t test, 2 tailed.

### Origins of Negritos and Aboriginal Australians

The Andamanese and the African pygmies seem obviously related in multiple aspects, including traits, Y relationship with the African megahaplogroup ABDE, and mtDNA haplotype M being closely related to African L. However, previous studies have found Andamanese to be even more genetically distant to Africans than other Eurasians [36]. Using the published genomes of 10 individuals from the Jarawa (JAR) and Onge (ONG) populations in the Andaman Islands [36], we found that Andamanese are relatively closer to Africans or have lower AFR/SAS(-BEB) distance ratio than other nearby populations such as BEB, with ONG more so than JAR, consistent with the known less admixture in ONG relative to JAR (Fig. 6A). PCA plots also showed Andamanese closer to Africans than all five populations of SAS (Fig. 6B). Relative to the distance to SAS, ONG showed smaller distance to Mbuti than to San or other Africans examined except LWK (Fig. 6C). The Mbuti group here consists of 4 published genomes from the Simons project [82] and the San group consists of 2 published genomes [93]. Given that Andamanese were closer to Africans than other Indians were (Fig. 6A) but Mbuti pygmies were not closer to Andamanese than some other Africans were, it can be inferred that Andamanese came from Mbuti rather than the opposite.

**Figure 6.**
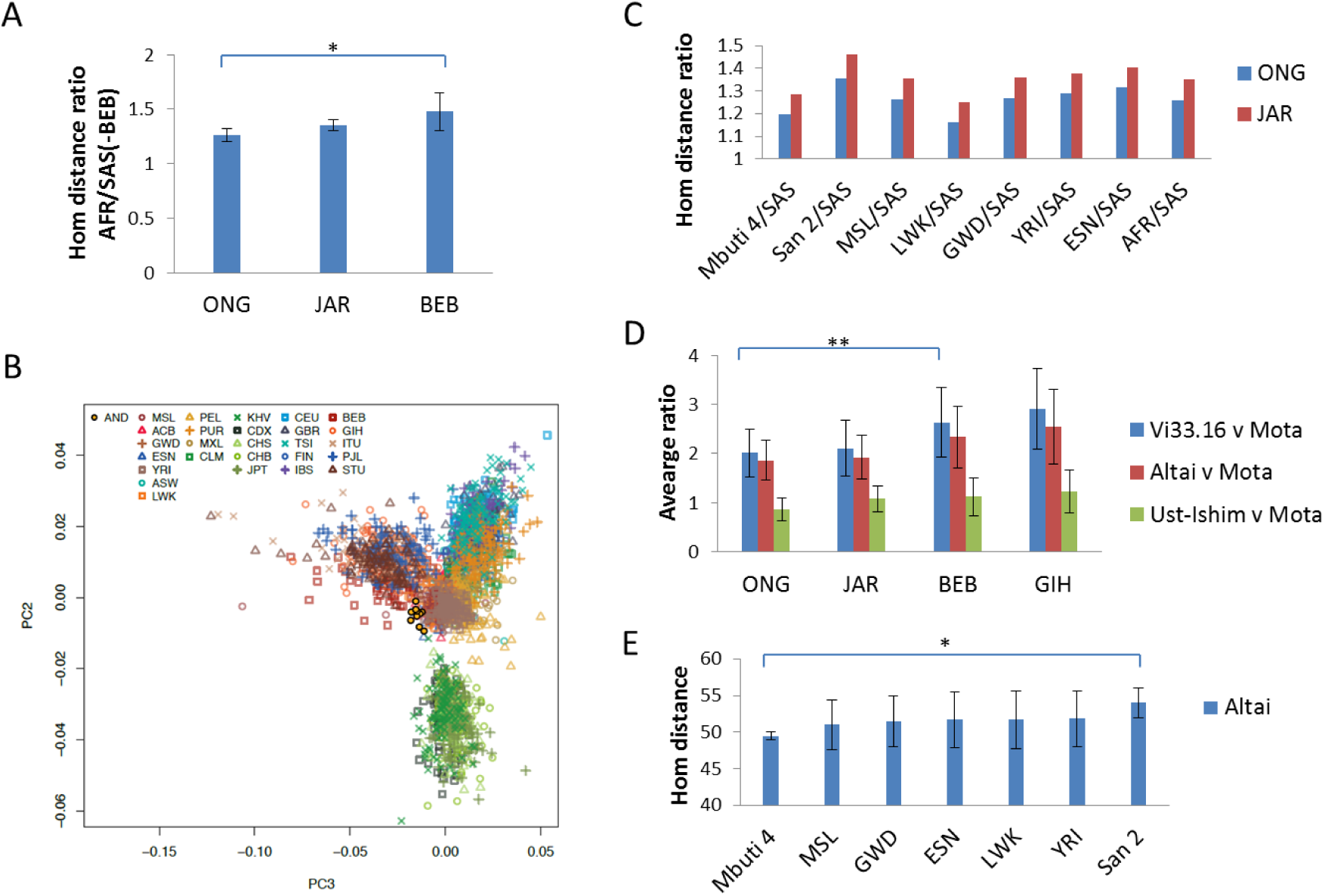
Origin of Negritos. (**a**) Shown are the ratios of ONG, JAR, or BEB autosomal distance to AFR versus SAS(-BEB). SAS (-BEB) excluded the BEB group from SAS groups. (**b**) PCA plot (PC3-PC2) analysis of 10 Andamanese and 1KG using slow autosomal SNPs. (**c**) Shown are the ratios of ONG or JAR autosomal distance to African groups versus SAS. (**d**) Hom distance ratio of ancient humans versus the Mota African for four South Asian groups (ONG, BEB, GIH, JAR). (**e**) Autosomal distance between Altai and various African groups. Data are means with standard deviation. **, P<0.01, t test or chi-squared test, 2 tailed.

The African affinity of Neanderthals prompted us to examine the distance between Neanderthals (with relatively higher coverage genomes, Vi33.16 and Altai) and several different Indian populations (ONG, JAR, BEB, and GIH) to see if ONG might have come from Neanderthals or related humans. Relative to the distance to the ∼4500 year old African Mota [94], ONG was closer to Neanderthals Vi33.16 and Altai, as well as to Ust’-Ishim who was known to have large amount of Neanderthal admixture, than other Indians were (Fig. 6D). Also, if Andamanese came from Neanderthals, Neanderthals should be closer to Mbuti than to San and other Africans, since Andamanese are closer to Mbuti than to San (Fig. 6C). This was indeed the case for the Altai individual who had high coverage genome for this analysis to be informative (Fig. 6E).

Since different Negrito groups in South Asia share similar traits, one expects them to be genetically related. The new Y tree grouping C with ABDE further suggests a common ancestry for different Negrito groups since the C haplotype is common in certain Negrito groups in Philippines while D is common in some others such as Onge. We therefore made use of a previously published SNPs genotyping data for a number of Oceanian groups including the Negrito group Mamanwa and its neighboring group Manobo in Philippines [63]. We measured the ONG/JAR distance ratio to look for the group that is closest to ONG relative to its neighbor JAR and the Mamanwa/Manobo distance ratio to look for the group closest to Mamanwa relative to its neighbor Manobo. Of the 13 groups examined, Mamanwa showed the smallest ONG/JAR distance ratio besides ONG; conversely, ONG showed the smallest Mamanwa/Manobo distance ratio besides Mamanwa (Supplementary Fig. S10). These results suggest that the two Negrito groups are more closely related to each other than either is to other groups as examined here.

We also examined the Aboriginal Australian (AUA) samples in the Pugach et al (2013) dataset and a previously published ∼100 year old AUA (AUA_100yr) who was unlikely to have admixed with European colonizers [95]. These AUA samples showed lower Mamanwa/Manobo ratio than other Oceanians (Supplementary Fig. S11). The AUA samples from Pugach et al (2013) also showed lower AFR/ASN ratio than other Oceanians, representing 68% of the average ratio for the Oceanians (excluding AUA and NGH or New Guinea Highlanders). To examine if the African component of AUA had come from Neanderthals, we calculated the Altai/ASN distance ratio of AUA and found it to be 64% of the average ratio for the Oceanians in Pugach et al (2013) dataset, which was significantly lower than the 68% found for AFR/ASN ratio, indicating closer relationship of AUA to Altai than to AFR. These results showed similarity between AUA and Negritos, indicating similar ancestry in Neanderthals and Denisovans.

### Testing the out of East Asia model

We next tested certain obvious predictions of the out of East Asia model. First, the model predicts lower diversity in people directly associated with the original AMH and higher diversity in people resulting from admixture of AMH with archaic humans. We calculated the hom PGD in slow SNPs as well as het numbers for each of the 25 groups totaling 2534 individuals in 1KG. The lowest hom PGD level was found in LWK followed by slightly higher level in CHS (Supplementary Fig. S12A). However, LWK has significantly higher numbers of het than CHS (Supplementary Fig. S12B). As high level heterozygosity indicates high genetic diversity and would reduce hom distance, it is likely that CHS has lower genetic diversity than LWK. We further found that within CHS (made of 72 individuals from Hunan and 36 from Fujian), Hunan samples have lower hom PGD and het numbers than Fujian samples (Supplementary Fig. S12CD). These results indicate that CHS, in particular Hunan people, have lowest genetic diversity levels among the 25 groups in 1KG. Given that known admixed groups such as MXL and PUR showed the highest genetic diversity or PGD (Supplementary Fig. S12A), it may be inferred that CHS or Hunan people may have the least amount of admixture and hence represent the original AMH group, at least among the 25 groups sampled here. That Africans, as human ancestor from ∼2 myr ago according to the multiregional model, did not show the highest genetic diversity level may seem unexpected but is in fact consistent with a key role for admixtures as claimed by the multiregional model as well as our out of East Asia model here. The original AMH group should have low admixture with archaic people in order for evolution into AMH to be possible since admixture may reverse AMH back to the archaic state.

Second, we would expect Southern East Asian groups to be closer to Africans. Although CHS represent samples collected from Southern China (Hunan and Fujian), while CHB (Han Chinese in Beijing) samples were from Northern China (Beijing), both in fact contain Southern and Northern Chinese. We therefore made use of the Hunan versus Fujian samples in CHS, where Fujian people are known to be mostly migrants from Central North China during the West Jin, Tang, and Song dynasties. We calculated the distance of each group to Hunan or Fujian and obtained the Hunan/Fujian distance ratio of each group. Consistently, groups known to have more Northern Chinese admixtures, such as CHB, MXL (Mexican Ancestry from Los Angeles), PEL (Peruvians from Lima), JPT (Japanese in Tokyo), had higher Hunan/Fujian distance ratio than Southern groups such as CDX (Chinese Dai in Xishuangbanna) and KHV (Kinh in Ho Chi Minh City, Vietnam) (Fig. 7A). Of note, FIN is closest to Hunan people among EUR groups (P<0.01), suggesting that North Western migrations of Southern Chinese during the first wave of AMH dispersal from Hunan area may have contributed to the ancestry of FIN. Consistently, Western hunter-gatherers from the Paleolithic age also showed closer distance to Hunan (manuscript in preparation). All AFR groups showed lower Hunan/Fujian distance ratio than non-Africans with LWK in East Africa the lowest, consistent with migration of Southern Chinese into Africa and into the Horn of Africa first. That non-Africans had more Fujian admixtures is consistent with known migrations of Northern East Asians into both the West and the America in more recent times during the Neolithic and Bronze ages. We further found Hunan people to be relatively closer to Africans than other South East Asians such as CDX and KHV (Fig. 7B), indicating origin of AMH more likely in Hunan relative to other nearby regions.

**Figure 7.**
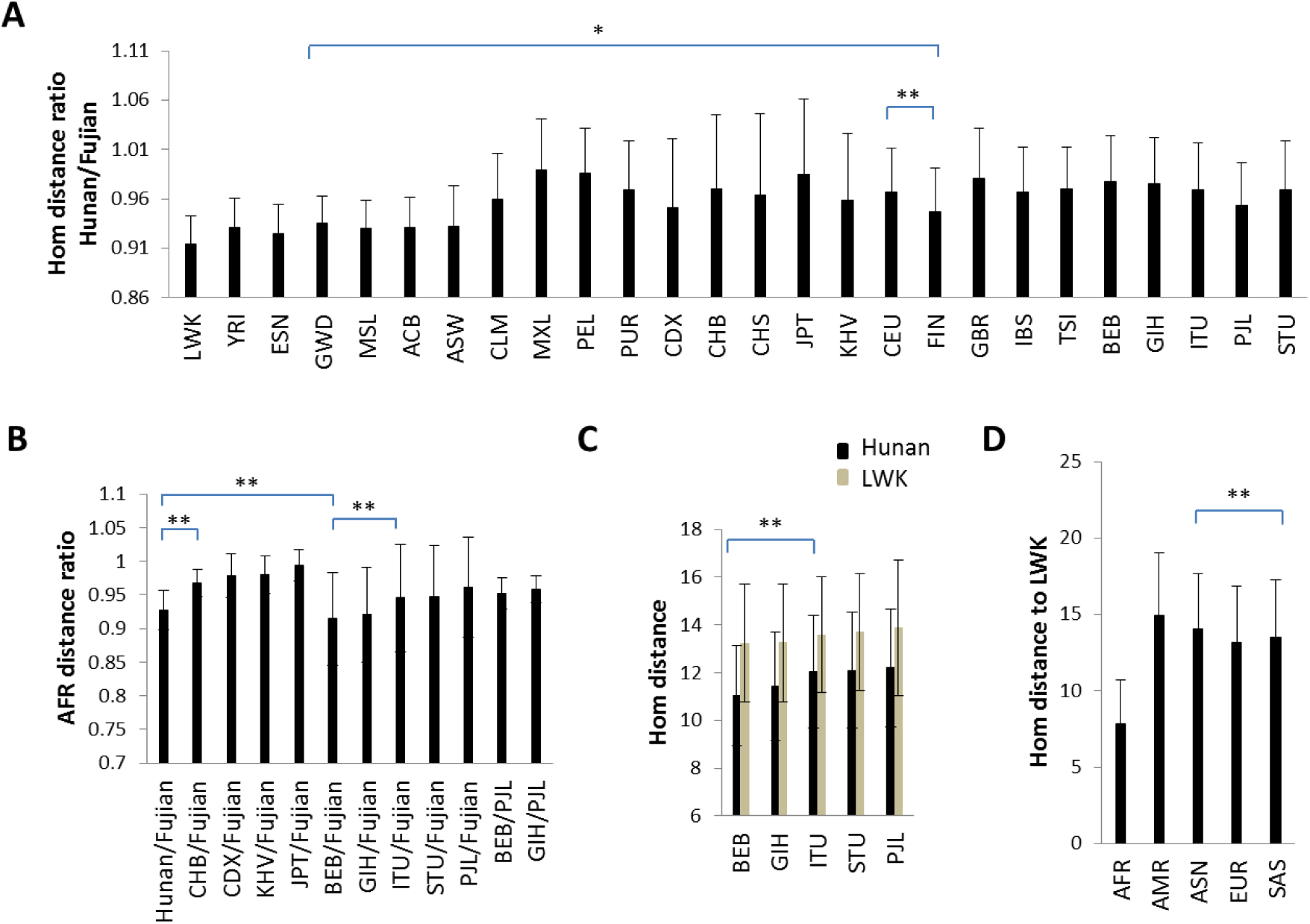
Hunan ancestry in Africans. (**a**) Ratios of autosomal distance to Hunan versus Fujian for each of the 25 groups in 1KG. (**b**) Ratios of autosomal distance to Hunan (or other East Asian and South Asian groups in 1KG) versus Fujian. (**c**) Autosomal distance to Hunan or LWK for various South Asian groups. (**d**) Autosomal distance to LWK for the 5 groups in 1KG. **, P<0.01, *, P<0.05, t test or chi-squared test, 2 tailed. Standard deviations are shown.

Third, as migration of AMH from Hunan via the Southern route to East Africa must cross the Indian subcontinent, one would expect closer relationship with Africans for groups within South Asia that are more related to Chinese relative to those more related to Europeans or more Southern relative to more Northern. Indeed, relative to Fujian people, the distance of different Indian groups to Africans follows exactly their direct distance to Hunan people, as well as their direct distance to LWK, in the order of increasing distance, BEB, GIH, ITU, STU, and PJL (Fig. 7BC). Also, Gujarati Indians (GIH) in Western India is closer to Africans than Punjabi people from Northern Pakistan (PJL) (Fig. 7B). Consistently, relative to PJL, both BEB and GIH are closer to Africans with BEB closer than GIH (Fig. 7B). The observation of lower BEB/Fujian distance ratio than Hunan/Fujian is consistent with Indians being in general closer to Africans than East Asians (Fig. 7D) and being more recent ancestors to Africans than East Asians based on the migration route of the out of East Asia model.

Fourth, we hypothesized that the branching process of Y may involve AMH hybridization with archaic humans and subsequent adaptive co-evolution of Y and admixed autosomes. As the first major split resulted in ABCDE, G, and HIJK haplogroups, we tested whether the ABCDE megahaplogroup, whose sub-branches are mostly found in Africans and South Asians or Oceanians with African like features, may have resulted from admixture of F AMH with admixed archaic Africans such as Neanderthals who may have migrated to South East Asia. We examined the Y chr sequences of three Neanderthals [6, 7], and found all to share alleles at informative sites with haplotype A0, A, AB and ABCDE but not with any non-ABCDE haplotypes (Table 3). These results further confirmed the African affinity of Neanderthals as shown by autosome analyses above and indicated that admixture of F AMH with Neanderthals may have resulted in African-like descendants with ABCDE megahaplotype who largely preferred to live in the Southern hemisphere. Consistently, East Asians (JPT) with D or C haplotype showed closer autosomal distance to Andamanese (also with D haplotype) or African MSL (with E haplotype) than those with O haplotype did (Supplementary Fig. S13).

**Table 3.**
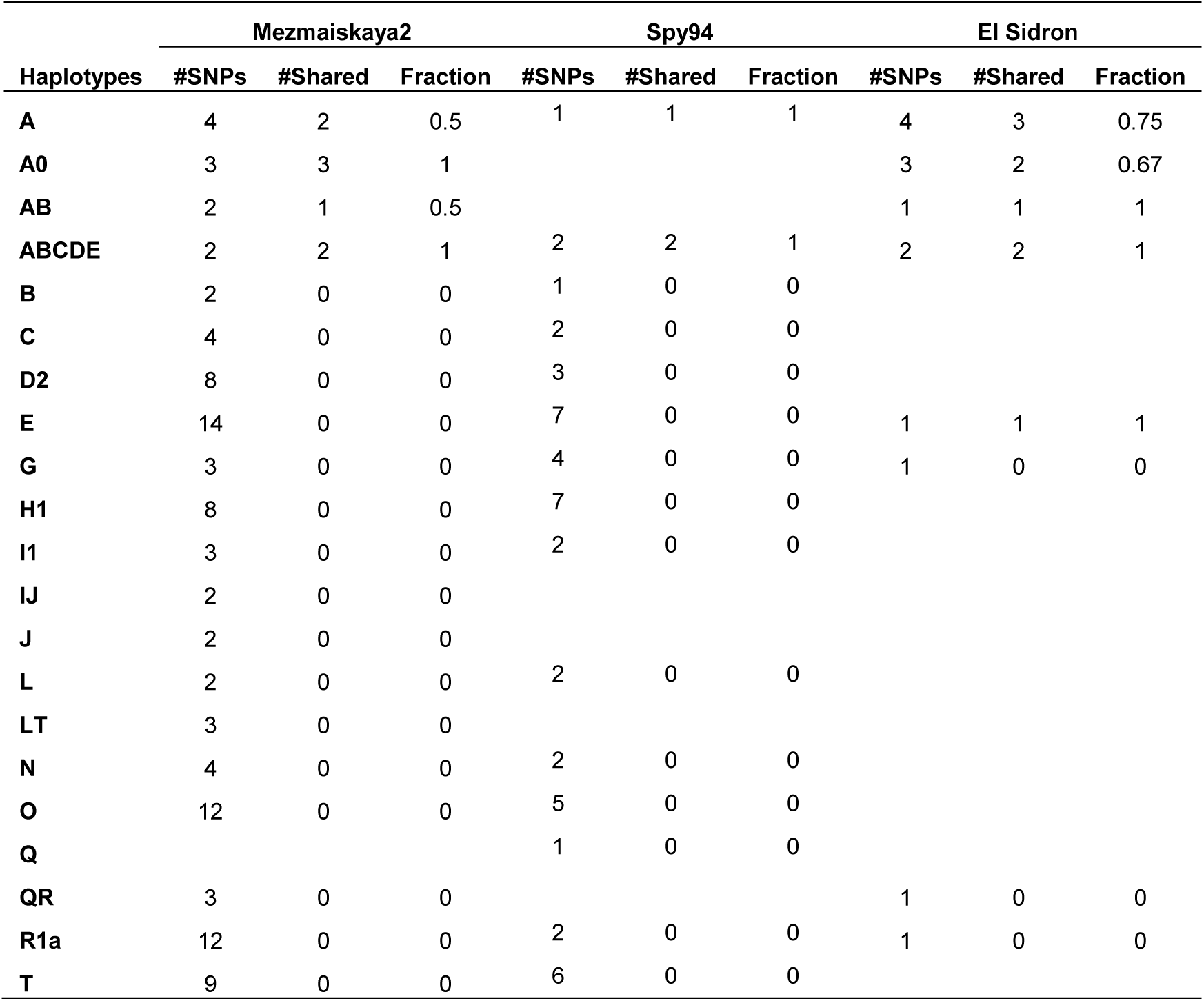
Sharing of Y chr alleles between Neanderthals and modern humans.

Fifth, to similarly test whether mtDNA diversification from the original ancestor type to more African type may involve AMH hybridization with archaic humans, we examined the distance between archaic and modern mtDNAs in slow SNPs (Supplementary Table S10). Although archaic mtDNAs were nearly equidistant as measured by mtDNA genome wide SNPs to the modern group consisting of Europeans (CEU: Utah Residents with Northern and Western European Ancestry), East Asians (CHS), and Africans (LWK: Luhya in Webuye, Kenya), they were closer to Africans in SNPs found in archaic humans (sites that differ between archaic mtDNA and the rCRS), indicating more sharing of archaic alleles in Africans (Supplementary Fig. S14). This is likely due to independent adaptive mutations since archaic mtDNAs are outgroups to modern mtDNAs as previous studies have shown. We also confirmed it by showing that the average distance between archaic and modern mtDNAs were larger than that within modern mtDNAs (Supplementary Fig. S15A). The archaic mtDNAs are at least of two types, with Neanderthal Vi33.16 and Altai belonging to one type or being close to each other than to other archaic mtDNAs while Denisovan and Heidelbergensis [96] belonging to another type (Supplementary Fig. S15B). Such results support the notion of multiple turnover events in mtDNA types in the past ∼2 myr of human evolution.

Our results here confirmed the first mtDNA tree ever built that placed the original AMH type (type 1 morph) in East Asia [29]. Our original type here is defined by the major alleles of 6 slow SNPs, 750, 1438, 2706, 8860, 14766, 15326 (Vi33.16 and Altai have all 6 except 2706, Denisovan has 14766 and 15326, and Heidelbergensis has 14766 and 1438). The earliest AMH Ust’-Ishim had the major alleles of these 6 SNPs and no additional slow SNPs. Mutation at 14766 defines V and VH and further mutation at 2706 defines H (Supplementary Fig. S5A). All other haplotypes overwhelmingly carry the major alleles of these 6 sites plus a few additional less common slow SNPs. We calculated the number of slow SNPs in each haplotype present in the 1KG and found R0 of Chinese to have the least amount among all non VH haplotypes, supporting R0 as the original type (Supplementary Fig. S16 and Table S11).

We examined whether the amount of allele sharing with archaic mtDNAs supports the above results linking archaic humans with South Asians or Oceanians. Using PhyloTree17 and Mitoweb data, we identified all non-L haplotypes that share alleles with archaic mtDNAs and calculated the number of shared alleles in each haplotype (Supplementary Table S12, Supplementary Information S4). Among R and N haplotypes, P4b1, R7a, J1b, and W3 were the most enriched with archaic alleles (4 alleles), which are common in Oceanians or Arabians/Caucasians/South Asians. Among M haplotypes (excluding L), G4, M17a, M27a, M76a, and M7b1a2a1a were the most enriched with archaic alleles (5 alleles), which are common in native Japanese, South East Asians, Papuans, and Indians. The M haplotype with the least amount of archaic alleles is D5 (1 allele), which is most common in Chinese.

We next calculated the distance of each modern haplotype in 1KG to archaic mtDNAs as measured by either slow or fast SNPs found in archaic genomes, and assigned each a distance ranking (Supplementary Fig. S17 and S18, Supplementary Table S13). Haplotype L0 commonly found in San people was the closest to archaic mtDNAs, consistent with the Neanderthal affinity with Y haplotype A. Haplotype G2 of JPT and M5a of SAS were top ranked non-L haplotypes in distance to Heidelbergensis in slow SNPs, consistent with the known admixed phenotypes of native Japanese and S. Asians. Relative to G2 that is common in South Asians and Tibetans, G1 is common in Russian far East and consistently closer to Altai in fast SNPs, indicating an adaptive role for fast SNPs. Consistent with the expected routes of human entry into America, D1 common in Amerindians and Paleoamericans is closest to Altai in both slow and fast SNPs among the 4 archaic mtDNAs. As expected, haplotypes common in African groups such as U6 and L3e are closer in slow SNPs to more African type archaic mtDNAs of Denisovan/Heidelbergensis, whereas those of Amerindians such as D1, D4 and A2 are closer to Neanderthals (Supplementary Table S14). Such analyses further indicate possible effects of archaic admixtures, with G2, I, T, and X2 affected by Denisovan/Heidelbergensis, and J, K2, and W by Neanderthals.

We next merged 1KG data with the AUA specific P4b1 and the Andamanese enriched M31a and found these haplotypes ranked among the top 13 in slow distance to Heidelbergensis (but 84th in fast SNPs), just following G and M5a among non-L haplotypes (Supplementary Fig. S19). Furthermore, they are uniquely much closer to Heiderlbergensis/Denisovan than to Altai, consistent with being uniquely related to Denisovan among living people today.

As M is defined by 10398G and 8701G, both present in archaic humans, it likely resulted from admixture of R0 with archaic Africans. While the effect of archaic humans can also be observed for some haplotypes within R and N, the M haplogroup may be the most affected as indicated by its defining SNPs and the extensive sharing of alleles between L (now within M) and archaic humans. Consistently, the ∼40000 year old Romanian Oase 1 had extensive Neanderthal admixture and carried an unusual N with the 8701G allele, indicating clear Neanderthal effect on its mtDNA [13]. The Oase 1 mtDNA may be an intermediate in the transition from N/R to M. A modern N haplotype with 8701G allele is N21 common in Malays.

## Discussion

We have arrived at a new model of modern human origins based on a more complete understanding of genetic diversity (Fig. 8). While the autosomes in our model are largely consistent with the multiregional hypothesis, the mtDNA and Y have a single origin in East Asia. We also identified Negritos and Aboriginal Australians as direct descendants of Neanderthals/Denisovans who were African migrants with Eurasian admixtures.

**Figure 8.**
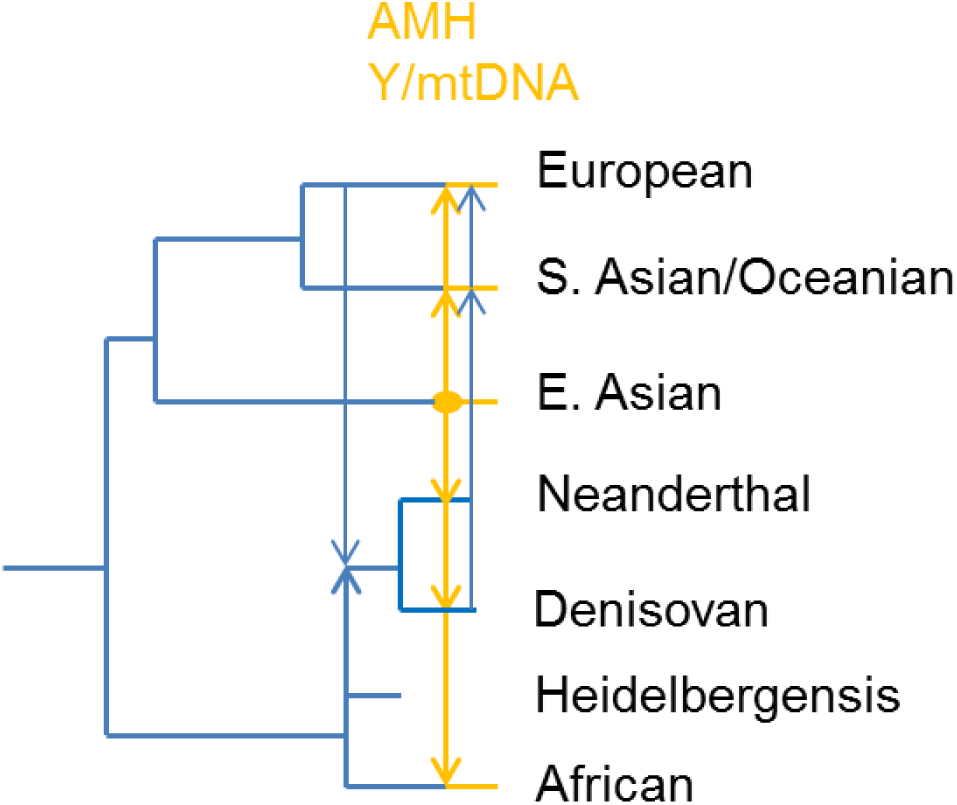
Model of human evolution. A schematic tree showing the phylogenetic relationship of major human groups, including Africans, East Asians, South Asians/Oceanians, Europeans, Heidelbergensis, Neanderthals, and Denisovans.

In highly conserved proteins, mutation rate may be inherently slower, as indicated by the lower mutation saturation relative to fast evolving proteins. The nonsyn SNPs in slow genes as defined here are neutral [62]. They are not deleterious and unlike the stop codon and splicing SNPs. They are also not under positive selection as positively selected genes tend to be fast evolving. The random set of SNPs, traditionally viewed as neutral, were here shown in fact to be similar to stop codon and splicing SNPs. Under the neutral theory, the slow and fast SNPs should produce the same phylogenetic tree topology. To the dramatic difference between slow and fast evolving DNAs as shown here, we cannot come up with a meaningful explanation using any known schemes other than the MGD theory [37].

Previous studies using genome wide SNPs (random or fast set) suggest a serial founder effect model resulting from the expansion of modern humans out of Africa, with intra-population diversity decreasing relative to their geographical distance from Africa and diversity between populations increasing with the geographical distances separating them [74–76]. However, such observations could be explained by ascertainment bias and different levels of saturation and selection pressures among groups. The finding here of more shared alleles among populations for fast versus slow SNPs suggests that the large fraction of shared alleles among populations is due to saturation and parallel mutations in fast SNPs, which has also been found in previous studies where higher sharing of alleles was found for microsatellite markers versus autosomal SNPs (microsatellites have higher mutation rates) [75].

We have shown that there are only three major human groups, Africans, East Asians, and Europeans/Indians. PCA plots using slow SNPs showed the first major division was between Africans and others (PC1), the second between East Asians and others (PC2), and third between South Asians and others (PC3). This supports the first split in humans to be between Africans and others, the second split between East Asians and others, and the third between South Asians and others. This indicates a Southern rather than Northern route for the first wave of AMH migration out of East Asia. The oldest AMH Ust’-Ishim clustered with South Asians. Also, the Y haplotype H of Indians diverged before diversification of European haplotypes, which is consistent with our model as well as the non-inhabitability for most parts of Europe during the Last Glacial Period. Aboriginal Australians and the related Negritos, traditionally viewed as the fourth major group, in fact consist of largely European/Indian and African genomes and their unique traits might have come from admixture of incoming Neanderthals with local archaic humans. Our calculation showed that the first major split of humans occurred 1.86-1.92 myr ago, well consistent with fossil evidence for the presence of *Homo* in Eurasia and the multiregional model. The coexistence at ∼1.76 myr ago in Africa of both Olduwan and Acheulean technologies suggests the coexistence of multiple groups of humans distinguished by separate stone-tool-making behaviors [97, 98]. The sudden appearance of Acheulean technologies and pro-Neanderthals at ∼0.5 myr ago in Europe (Sima de los Huesos site of Atapuerca) can now be explained by a more recent out of Africa migration by the ancestors of Mbuti people [99, 100].

Mitochondrial DNA (mtDNA) and the non-recombination region of Y chr (NRY) lack recombination and provide records of history that are independent of autosomes. Most SNPs in these DNAs can be proven to be under selection, e.g. certain SNPs or haplotypes of mtDNA or Y chr are known to be related to human diseases or compatibility with nuclear genomes [43, 101–105]. Sharing of alleles of mtDNA or Y chr should mean similar selection, reflecting both environments and physiology or primarily physiology when saturation has been reached. Sharing of physiology should be informative for a phonetic approach of phylogeny. Coevolution of mtDNA, Y, and autosomes has been found by many studies [43, 105–108], which may play a key role in the diversification into multiple haplotypes during AMH radiation from its place of origin to other regions by hybridization with archaic humans. People who have stayed relatively unchanged in physiology and living environments from the ancestor would be expected to have few deviations from the ancestor haplotype and their present day living place would indicate place of origin for the ancestor. It is through such reasoning that we have come to place the origin of modern Y and mtDNA in East Asia or Southern China. Our results showed that groups with the same Y or mtDNA haplotypes are also closer in autosomes and traits. The Y megahaplogroup ABCDE matches with the mtDNA megahaplogroup M. Such *a priori* sensible results provide strong independent validation for our new phylogenetic method.

Given that most SNPs in Y and mtDNA are not neutral, one cannot use the molecular clock approach to determine the age of the haplotypes except for recent diversifications. That the Y haplotype NO of the ∼45,000 year old Ust’-Ishim differs from the putative ancestor F by only ∼27 SNPs whereas a present day haplotype could differ from the F ancestor by as much as ∼740 SNPs (Fig. 3) indicates that the ancestor F should not be much older than ∼45,000 years. This relatively young age is remarkably consistent with the time point for the replacement of Neanderthals by AMH but appears to contradict the oldest AMH fossils in Africa or in Hunan China [21]. However, nearly all AMH fossils older than 40,000 years still have certain archaic features and independent evolution of modern features has been noted to occur periodically over the past 950,000 years since the time of *H. antecessor*[4, 109].

The novel concept here of modern replacing archaic versions of Y and mtDNA but not autosomes is key to our model of out of East Asia. The lack of recombination in Y and mtDNA makes this idea biologically inevitable. The fact that Heidelbergensis, Denisovans, Neanderthals, and AMH all have distinct mtDNAs suggests that such replacements may have taken place multiple times in the past. Modern examples consistent with the replacement idea are the dominant presence of Asian Z mtDNA in the Saami people of Northern Europe and the wide presence of Asian Y haplotype N in Finnish, who are otherwise largely indistinguishable from Europeans in both autosomes and traits. Also consistent is the finding of three super-grandfather Y haplotypes in China that are relatively young in age (∼5000-7000 years) but account for ∼40% of Han Chinese males today [110, 111]. Admixture of incoming Asian AMH with archaic humans in Europe or Africa would lead to haplotype diversification in Y and mtDNA while still maintaining regional specificity in autosomes and hence traits as traits are mostly determined by autosomes. Therefore, the multiregional model is fully compatible with any single origin model of mtDNA or Y chr and there is no real conflict between the timing of autosome diversification and the much more recent appearance of the modern mtDNA and Y. Interbreeding between modern and archaic humans may best explain certain human fossils near or within the era of AMH that show both archaic and modern features, such as *H. naledi* and the „Red Deer Cave‟ people [112, 113].

The Out of Africa model requires uni-parental DNAs and autosomes to be both modern to qualify as AMH but our model only requires uni-parental DNAs. The Y tree topology shows features not consistent with the Out of Africa model. The Y tree shows haplotype I splitting before R from NO and so I should carry less Asian autosomes than R if the Out of Africa model is true but not if our model is. But, Europeans carrying I and R are not known to be different in autosomes. Also, the Out of Africa model has divergence of Europeans and Asians in the Middle East. However, the Y tree shows West Asia haplotype J splitting after the differentiation of South Asia haplotype G, indicating differentiation of Eurasia types following type F to be in East Asia rather than West Asia, which is unexpected by the existing model. Recent ancient mtDNA studies found the earliest modern R to be close to the earliest fossil date of AMH at 45 ky ago and 5000 years older than the earliest N, thus invalidating the existing tree and validating our Out of East Asia tree here [114].

The ∼45,000-year-old AMH Ust’-Ishim from Siberia was previously found to have left no descendants among present populations and to be more related to East Asians than to Europeans/Indians [14]. However, our results showed this individual as Indians. This discrepancy is to be expected. It has been routinely found as surprising in previous studies on ancient DNAs that there is no genetic continuity between ancient and present day people. Such unexpected anomalies can now be understood as artifacts of using non-informative or fast SNPs which turned over quickly to be adaptive. We have verified this for most ancient European DNAs that while non-informative SNPs placed them as outliers, the slow SNPs as defined here all placed them as indistinguishable from present day Europeans.

Our finding of Neanderthals and Denisovans as primarily Africans with Eurasian admixture is well supported by fossil data indicating *H. heidelbergensis*, present in both Africa and Europe, as ancestors of Neanderthals. The taurodont teeth are common in Neanderthals, Heidelbergensis and certain South African fossils [115]. The occipital bunning of Neanderthals are also common in modern Africans [116]. Neanderthals are known to share multiple traits with Europeans such as the prominent shape and size of the nose [35, 117], which supports our finding that Neanderthals were often closer to Europeans than to E. Asians (8/10 examined here). Thus, traits and genotypes are coupled after all as they should, unlike the strange and surprising conclusion of more Neanderthal alleles in East Asians than in Europeans as analyzed by using non-informative fast SNPs [11, 12]. Our result that Denisovan is nearly equally related to East Asians and Europeans (slightly more related to East Asians) is consistent with where Denisovan was found. Seemingly unexpectedly, certain Neanderthals found in Europe was most closely related to Asians (Vi33.25) or Americans (Vi33.26). However, this would be expected if Africans associated with the Neanderthal exit had entered Asia or South Asia via the Northern route from Siberia or possibly a Southern route. The general lack of Neanderthal fossils in this Southern route may reflect the relatively small effort so far invested in this region (with only few *Homo* fossil finds like Narmada from ∼200 kya who is broadly classified as *H. heidelbergensis*). Indeed several fossils in China show Neanderthal features such as the inner-ear formation in the ∼100 ky old *Xujiayao* and *Xuchang Man* [4, 118–120]. Certain mysterious Southern China fossils such as the 11-15.5 ky old „Red Deer Cave‟ people with hybrid features of modern and archaic humans may also be candidates for Asian relatives of Neanderthals, especially considering their taurodont teeth [112]. Early modern human fossils with typical Mongoloid features in South West China (Liujiang, Ziyang, Lijiang, and Chuandong) also have weak occipital buns commonly found in Neanderthals [4, 119, 121]. Thus, although Neanderthals were mostly found in Europe and Middle East, they likely also made their way to North East Asia (Denisovan and Teshik-Tash) and South East Asia [122].

Fossils or traits indicating AMH migration from East Asia into Africa or Europe have been noted before. First, native Africans such as Khoisans are well known to have certain East Asian features such as shoveling teeth, epicanthic fold, and lighter skins. Mbuti pygmies look very much like the Andamanese. The much lower frequency of shoveling teeth in African fossils and Khoisan relative to ancient and modern Chinese suggests that this type of teeth could only originate in China with its African presence due to migration. The type of shoveling teeth found in Neanderthals and Pleistocene *Homo* from Atapuerca-Sima de los Huesos may either be a different type from that of Asians and Africans or come from early disposal of *Homo* from Asia to Europe [123, 124]. Second, a combination of three features has been noted to be region-specific to China fossils with lower frequency also found in North Africa: a non-depressed nasal root, non-projecting perpendicularly oriented nasal bones and facial flatness [125]. Third, Dali man of China (∼260 kya) had lower upper facial index and flat nasomolar angle, but these two modern features only first appeared in Europe in Cro Magnons (Xinzhi Wu, personal communication).

The appearance of modern humans should be accompanied by new technologies just as the knife type stone tools were associated with the first appearance of the genus *Homo*. A technology just one step more advanced than stone tools is pottery making. Consistent with our model, the earliest pottery making intended for practical usage was found in Hunan and the neighboring Jiangxi in Southern China at 18,000-20,000 years ago [126, 127]. While future investigations could extend the time even earlier, one should not expect a new technology to appear simultaneously with the first appearance of AMH since it would take time for the first modern humans to grow into a large enough population to be able to invent new cultures. It is also remarkable to note that the next new invention after pottery, rice or agriculture, also likely came from Hunan [128]. Hunan is also the site of the earliest fully modern human fossils [21]. Placing AMH origin in China is also in line with the observation that the best argument for regional continuity has been built using data from China [4]. The observation here that different modern Chinese people could have independent genetic lineages separated by hundreds of thousands of years is consistent with the morphological observation that *H. erectus* and *H. sapiens* in Northern China are not identical to those in Southern China [129]. Among all East Asians examined here, the genomes of Hunan people were found most enriched in Africans. Therefore, our model of modern human origins in East Asia, in particular Hunan Province in China, provides a satisfying account of all relevant data.

The study here shows different genetic diversity levels in different human groups depending on different types of SNPs. Europeans show the lowest genetic diversity level in stop codon and splicing SNPs while Africans the highest, which has also been found in a recent study [130]. However, East Asians show the lowest genetic diversity in genome average and hence in non-coding sequences. Thus, different populations encounter different selective pressures, the precise nature of which would require future research. Already, however, some tentative hints emerge on the genetic basis of certain complex traits that are commonly thought to be culturally shaped. The difference in selective pressure on non-coding or regulatory regions versus proteins or parts is reminiscent of the thinking style difference between the East and West in philosophy and medicine, i.e., the holistic versus the analytical [131]. The high genetic diversity of Africans may confer adaptive advantages as in high diversity in immunity [132].

## Conclusions

The MGD theory provides a more complete understanding of the long standing puzzle of what determines genetic diversity, which makes inferring human origins from genetic diversity patterns realistically possible. By better identification of phylogenetically informative genes and constraining neutral theory application to these genes, we provide strong molecular evidence for multiregional evolution of autosomes and for East Asia origin of modern Y and mtDNA. Further work utilizing the MGD theory is ongoing and may yield more surprising and yet satisfying results in human evolution.

## Abbreviations

AMH: anatomically modern humans
MGD: maximum genetic diversity
SNP: single nucleotide polymorphisms
AUA: Aboriginal Australian
PGD: pairwise genetic distance
PCA: principal component analysis
Myr: million years
AFR: African
ASN: East Asian
EUR: European
SAS: South Asian
ESN: Esen in Nigeria
GBR: British in England and Scotland
CHS: Southern Han Chinese
CHB: Han Chinese in Beijing
JPT: Japanese in Tokyo
BEB: Bengali from Bangladesh
YRI: Yoruba in Ibadan, Nigeria
CEU: Utah Residents with Northern and Western European Ancestry
LWK: Luhya in Webuye, Kenya

## Acknowledgements

We thank Shuhua Xu, Xitong Lu, Denghui Luo, Jie Liang and Xiaohua Tan for technical assistance. We thank Xinzhi Wu for sharing unpublished work. We are grateful to Mark Stoneking, Irina Pugach, David Reich, Joseph Pickrell, Arti Tandon, Sarah Tishkoff, Joseph Lachance, Philip Johnson, and Brenna Henn for sharing DNA datasets. Although not all analyses of these datasets were presented here, they have been helpful to make this work possible. We thank Xinzhi Wu, German Dziebel, and Joseph Daniels for critical reading of the manuscript. We thank Feng Gao, Xing Gao, Yamei Hou, Wu Liu, Erik Trinkaus, Lingxia Zhao, and Changqing Zeng for valuable discussions.

## Ethics approval and consent to participate

Not applicable.

## Consent for publication

Not applicable.

## Availability of data and material

The datasets generated and analyzed for this study are available in the Additional files.

## Author contributions

SH and DY designed the study. DW and JY identified the slow evolving proteins. DY, XL, YG, MW, YZ, ZZ, and SH performed data analyses. SH and DY wrote the manuscript and all authors made comments.

